# Nanometric axial localization of single fluorescent molecules with modulated excitation

**DOI:** 10.1101/865865

**Authors:** Pierre Jouchet, Clément Cabriel, Nicolas Bourg, Marion Bardou, Christian Poüs, Emmanuel Fort, Sandrine Lévêque-Fort

## Abstract

Strategies have been developed in LIDAR to perform distance measurements for non-coherent emission in sparse samples based on excitation modulation. Super-resolution fluorescence microscopy is also striving to perform axial localization but through entirely different approaches. Here we revisit the amplitude modulated LIDAR approach to reach nanometric localization precision and we successfully adapt it to bring distinct advantages to super-resolution microscopy. The excitation pattern is performed by interference enabling the decoupling between spatial and time modulation. The localization of a single emitter is performed by measuring the relative phase of its linear fluorescent response to the known shifting excitation field. Taking advantage of a tilted interfering configuration, we obtain a typical axial localization precision of 7.5 nm over the entire field of view and the axial capture range, without compromising on the acquisition time, the emitter density or the lateral localization precision. The interfering pattern being robust to optical aberrations, this modulated localization (ModLoc) strategy is particularly well suited for observations deep in the samples. Images performed on various biological samples show that the localization precision remains nearly constant up to several micrometers.

---

In the presence of coherent signals, interferometry offers unmatched sensitivity for distance measurements^1^. Measuring the relative phase between the excitation in the elastically scattered or reflected signal reaches record precisions. Interferometry has thus naturally been used in some coherent microscopy configurations to obtain nanometric axial localization^2–5^. However, in the case of fluorescence microscopy which is today the most widespread technique for cell imaging such an approach remains impossible because of the non-coherent fluorescence emission with respect to the illumination. In single molecule localization microscopy (SMLM)^6–9^ for which axial localization precision needs nanometric precisions, interference based strategies would really be beneficial.

Effectively, axial localization is still an ongoing central issue in SMLM^10^. The symmetry of the microscope and the optical optimization of the objectives in the lateral direction deteriorate the axial localization precision compared to the lateral one. A large variety of strategies have been developed based on point spread function (PSF) manipulations like multi-focused techniques^11–14^, PSF engineering approaches^15–18^ or supercritical angle fluorescence detection^19–21^. Alternative accurate techniques are based on the coherence of the fluorescence when emitted by a single molecule to retrieve axial localization, like interferometric PALM (iPALM)^22,23^, 4Pi single marker switching nanoscopy (4Pi-SMSN)^24,25^ or Self-Interference (SelFi)^26^.

The axial localization is often performed at the expense of the lateral one, resulting in a reduced optimized working depth^18,27^ or field of view^26^. Most techniques are optimal in the focal plane, and quickly degrade with defocus. In addition, their performances are hindered by optical aberrations within the sample beyond a few micron depths unless adaptive optics^25,28,29^or more robust fluorescence self-interference approaches^26^ are used.

Distance measurements is also being intensively addressed by the optical community developing Light detection and ranging (LIDAR) systems^30,31^. The distance is typically retrieved by time of flight (ToF) measurements of the light, from the source to the target and return. The most widespread technique used in vehicles and video game applications is called amplitude modulated continuous wave LIDAR ^32^. This LIDAR strategy smartly bypasses the non-coherent detection by introducing an alternative amplitude modulation in the excitation beam. Since the targets respond linearly to the excitation intensity, the distance is measured from the relative phase between the detected signal and the reference modulation. This latter is usually performed by electronically modulating the laser amplitude at high frequencies, typically up to 100 MHz^33,34^. In the case of sparse point-like targets the phase localization precision turns out to be much more accurate than the spatial period of the excitation (>1 m) but since this latter is much larger than the excitation wavelength the resulting localization precisions fall in the millimeter range. Here we propose to use a modified configuration of the amplitude modulated LIDAR strategy to perform nanometric axial localization of single fluorescent molecule in the context of SMLM. To achieve such a precision, the excitation modulation is performed by shifting interferences with a tilted light beam. This results in spatial period of the excitation of the order of the wavelength and in the decoupling of the time and spatial modulation frequencies. By acting as local probes in a known excitation field, single molecule positions are revealed directly by the phase of their modulated emission. This new axial modulation localization (ModLoc) method can operate with a standard single objective configuration. We show that the precision remains uniform over the capture range and is robust to optical aberrations, ModLoc thus enables imaging up to several tens of microns inside the biological samples.

## Principle

Figure 1a shows the principle of a standard amplitude modulated continuous wave LIDAR. The intensity of a continuous monochromatic laser source at the optical angular frequency *ω* and excitation wavelength *λ*_*exc*_ is modulated by an electronic modulator at the angular frequency Ω ≪ *ω*. The resulting spatial period of the modulated excitation beam is given by 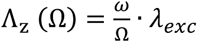. Since the response of the target is linear in intensity, the signal is also modulated at Ω. The source/target distance *z* is obtained by a lock-in detection which measure the phase shift *ϕ* between the local oscillator and the detected signal. It is given by *z* = (*λ*_*exc*_/2*π*)(*ω*/Ω)*ϕ* (modulo Λ_z_(Ω) due to phase wrapping). The localization precision depends on the precision of the phase measurement. The presence of the huge coefficient *ω*/Ω (typically > 10^6^) strongly impairs the localization precision and impedes its use in microscopy. Figure 1b shows the proposed alternative implementation in which the spatial modulation of the excitation beam is performed by adding an interfering tilted beam. This configuration, designated as ModLoc in the following, offers a versatile control of the illumination pattern: the angle α between the two beams determines the fringe spatial period Λ and the angle *θ* between their bisecting line and the coverslip axis is equal to the angle of the fringes with the optical axis (see Supplementary Fig. 3, in Fig. 1b *θ* = *α*/2).

**Figure 1.**
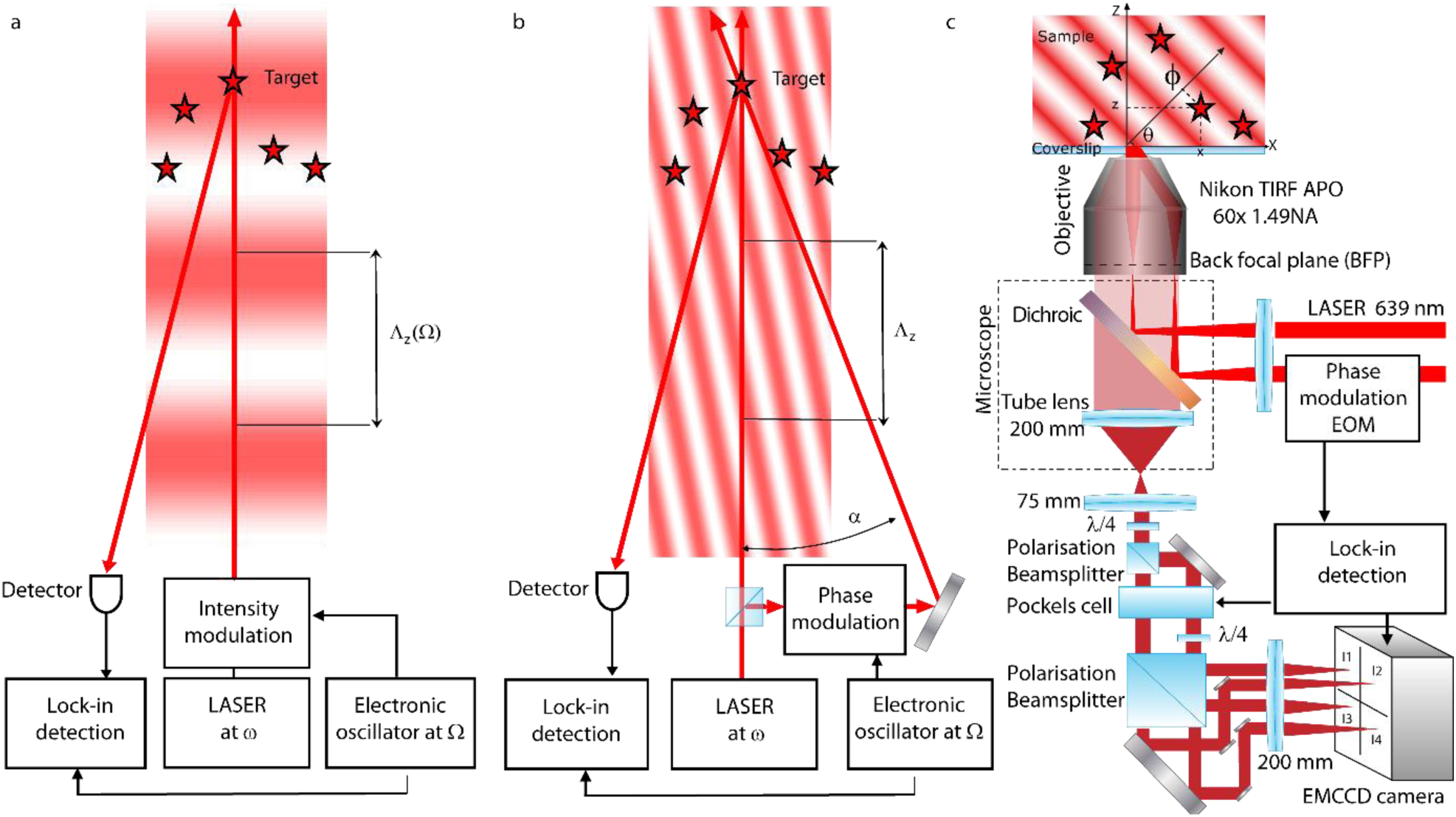
Modulated localization (ModLoc) principle and experimental implementation. **a**, principle of a standard lock-in amplitude modulated continuous wave LIDAR for which the excitation modulation is performed by electronically modulating the laser intensity at the frequency Ω. **b**, Alternative ModLoc implementation for which the excitation beam spatial modulation is performed using an interfering tilted beam with an angle *α*, the spatial and temporal modulations are fully decoupled and can be set independently. The ModLoc configuration introduces an additional phase shift which depends on the transverse position *x* of the target due to the tilted interference pattern. **c**, Implementation of ModLoc microscopy: The tilted illumination pattern (*λ*_*exc*_ = 639 nm) is obtained by the interference of two parallel beams focused in the back focal plane of the 1.49 NA objective with tunable off-axis distance to control the interference period and the fringe angle *θ* in the sample. The time modulation frequency is performed by shifting the relative phase between the two excitation beams using an electro-optic modulator (EOM). The lock-in demodulation is performed thanks to a dedicated detection module: a *λ*/*4* plate is followed by a polarization beamsplitter, a Pockels cell, a *λ*/*4* plate in one of the path, and a last polarization beamsplitter. The initial fluorescence image is steered into a set of 4 detection channels with complementary time varying transmission performed by a Pockels cell and synchronized with the EOM. These 4 channels are imaged simultaneously on the EMCCD camera. The modulation/demodulation frequency, typically 600 Hz, is chosen to be compatible with the short ON-time of the emitters averaging over typically 30 modulation cycles for each acquired EMCCD frame. For details see Supplementary Fig. 1-3 and Note 1-2)

In addition, this configuration enables a complete decoupling between the spatial and temporal modulations which can be set independently. The time modulation is obtained by shifting the relative optical phases between the two excitation beams. The source/target distance *z* = Λ_z_(α, θ)*ϕ*/2*π* is retrieved using a standard lock-in detection.

The ModLoc configuration introduces an additional phase shift which depends on the transverse position *x* of the target due to the tilted interference pattern (see Fig. 1b) given by:

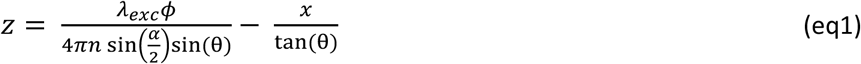

The axial localization precision is thus given by the sum of two terms: a phase localization term and a lateral localization term typically given in SMLM by centroid fitting of the PSF. The axial standard deviation *σ*_*ModLoc*_ can be written as:

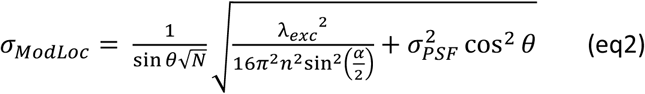

with *N* being the number of detected photons involved in the phase detection and *σ*_*PSF*_ the PSF size (see Supplementary Note 3). The axial modulation period Λ_z_(α, θ) is of the order of *λ*_*exc*_ for large angles α and *θ*. These angles are readily accessible with the high Numerical Aperture (NA) objectives used in SMLM. The ultimate limit for a single objective excitation configuration, albeit for grazing angles, is given by *θ* = *α*/2 = 45° and results in a quasi-isoprecision of *σ*_*ModLoc*_ ≈ 1.1 *σ*_*x*_, *σ*_*x*_ being the standard deviation in the lateral direction *x*.

For SMLM, the demodulation frequency range must be compatible with the ON-time properties of the emitters. This ON-time follows a Poissonian statistic with an average value *τ*_ON_ which depends on the illumination intensity and the chemical properties of the molecule and its environment.^35,36^ The modulation frequency must satisfy 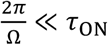 to retrieve the phase for the large majority of the emitters. In addition to fulfill time sparsity and localize most of the molecules in a single shot, *τ*_ON_ must satisfy *τ*_ON_ < *τ*_cam_, *τ*_cam_ being the minimum exposure time of the camera to optimize the acquisition time. Thus, the demodulation must be faster than the acquisition rate and needs a specific optical system to perform the lock-in detection.

Figure 1c presents the experimental implementation of the ModLoc microscope (Supplementary Fig. 1 and Note 1 for details). The tilted illumination pattern is obtained by the interference of two parallel beams focused in the back focal plane of the high NA objective: the off-axis distance of each beam is tunable independently to control the interference spatial period and the fringe angle *θ* in the sample. The time modulation frequency is performed by shifting the relative phase between the two excitation beams using an electro-optic modulator (EOM). The lock-in demodulation is performed thanks to a dedicated detection module based on a Pockels cell which sorts the initial fluorescence image into a set of 4 detection channels with complementary time varying transmission synchronized with the excitation modulation (Supplementary Fig. 2 and Note 2 for details). Each channel is imaged into a specific subarray of the camera. Several transmission functions are possible for the channel sorting, but they should be complementary to avoid any photon loss. The time integration is performed by the camera over several modulation periods. The full decoupling between the spatial and temporal modulations in ModLoc gives a necessary degree of freedom for SMLM applications. Typically, the modulation frequency is set to 600 Hz for an acquisition rate of the camera of 20 Hz with 30 modulation cycles for each frame. As for classical SMLM imaging an average of 10000 to 20000 images are typically acquired to reconstruct the final super-resolved image.

Thanks to sparsity, the fluorescence intensity *I*_*q*_ transmitted into the channel *q* by each fluorophore after integration by the camera can be measured. Its phase associated to the ModLoc detection is obtained readily by 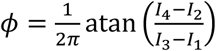 and its axial position *z* can be deduced. To avoid phase wrapping, Λ_z_(*α*, *θ*) is set to be larger than the depth of focus, approximately 1 μm (Supplementary Fig. 5 and Notes 4). The optimized axial localization is obtained with α = 42° and *θ* = 44°. In this configuration with the ModLoc implementation (Fig. 1c), the expected axial standard deviation satisfies *σ*_*ModLoc*_ ≈ 2.5 *σ*_*x*_ (see Supplementary Note 3) and can be used to image from the coverslip to several tens of microns in depth.

## Results

We first characterize ModLoc axial and lateral localization precisions at various depths. To evaluate the intrinsic precision, we perform repeated localizations over small fluorescent nanospheres with a controlled emission level as previously reported^21,22,26^ (see Supplementary Note 5). Figure 2a shows the very good agreement between the experimental data and the theoretical limits (Cramèr Rao Lower Bound, CRLB) for various levels of photons. For a typical number of photons of Alexa 647 emitters (~3500 photons), ModLoc achieves an axial localization precision of about 7.5 nm. Figure 2b shows that the axial and lateral localization precision over the whole capture range remains constant over the whole capture range. In addition, unlike most PSF shaping methods, the lateral precision is not compromised on the axial one. The accuracy, representing the discrepancy between the measured position and the real one, is another important parameter in SMLM. This parameter is evaluated from images of the bottom part of fluorescently labeled (AF647) 3µm diameter microspheres (Figure 2c) and their associated xz projection (Figure. 2d, Supplementary Note 5). The measured positions of different localized fluorescent probes coincide with their theoretical positions on the sphere (Figure 2e). The performances of ModLoc in depth are evidenced by repeated localizations of small nanospheres for various axial positions up to 7 µm. Figure 2f shows that the lateral and the axial localization precisions are constant over this extended depth and equal to respectively 3.3 nm and 7.5 nm. These findings prove that ModLoc is a real asset for 3D single molecule imaging especially for in depth imaging.

**Figure 2.**
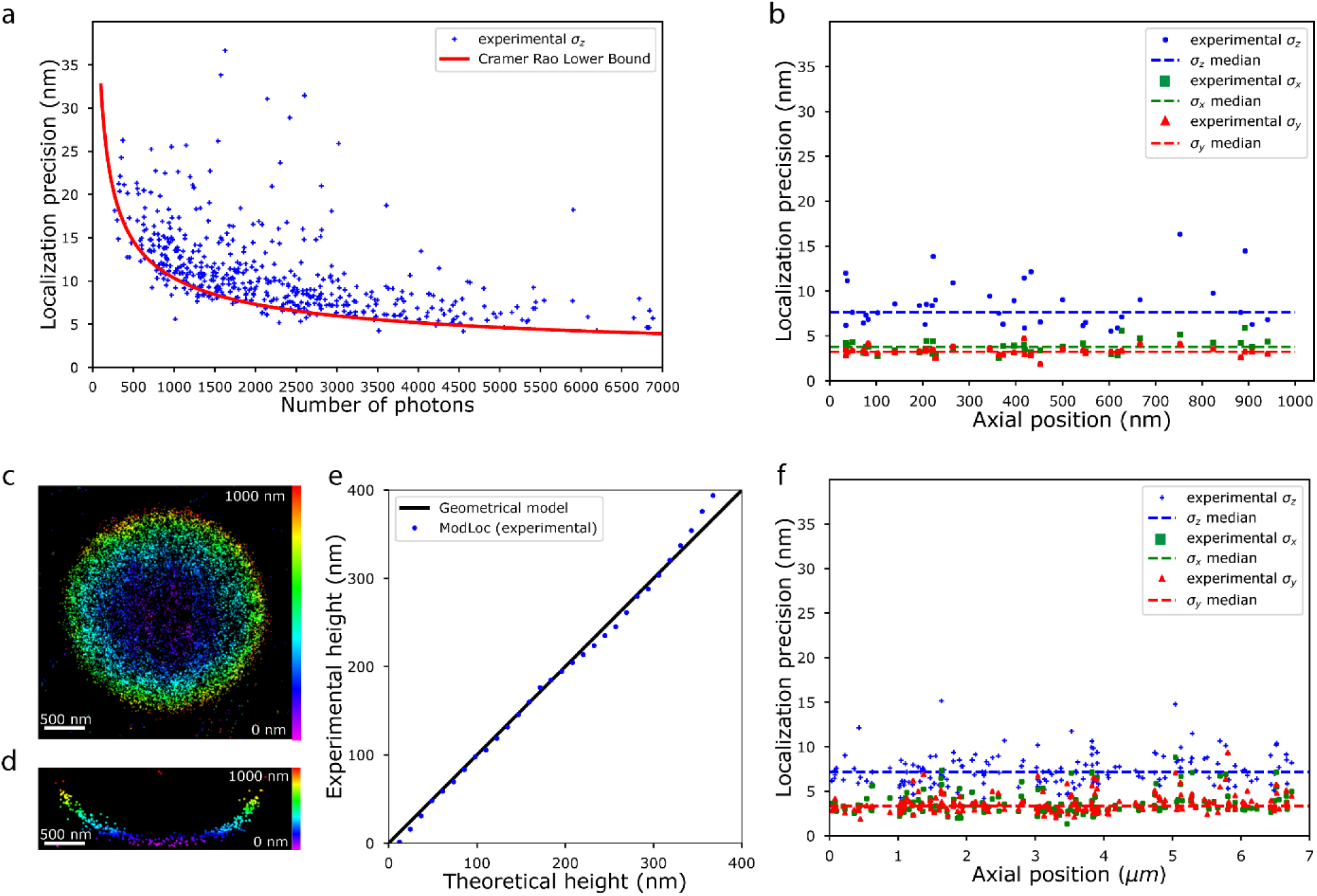
Performances of ModLoc. **a**, Theoretical and experimental localization precision of ModLoc as a function of the number of photons. The experimental localization precisions are obtained from the standard deviation of the localization distribution for different 40 nm diameter red fluorescent beads. The measurement for all different beads was done over 50 frames, the red curve represents the Cramèr-Rao lower bounds. **b**, Comparison between lateral (green and red dots) and axial (blue dots) localization precision over the whole capture range in the first micron. Results are extracted from previous representation **a** by keeping results comprise between 2500 and 6500 photons and an effective pattern contrast superior to 0.6. **c**, 3D super resolved image of a 3 micrometers diameter sphere labelled with AF647 fluorescent probes conjugated to streptavidin, the capture range limits the observation to the bottom of the sphere close to the coverslip. **d**, Sphere xz cross section shows the geometric profile of the micro-sphere. **e**, Comparison of the experimental heights of different single molecules detected and the theoretical heights calculated from the sphere geometrical model. The black curve represents the expected tendency and median values of ModLoc are represented by blue dots. **f**, ModLoc axial localization precision and lateral precision at different depths up to 7 µm. Repeated measurements (50 Images) were made on 40 nm diameter red fluorescent beads deposited on non-labelled cells (see supplementary note 5 for more details). The number of photons are between 2500 and 6000 and the effective contrast measured is superior to 0.6.

We now focus on ModLoc performances on biological structures (Supplementary Note 6 and 7). COS7 cells are imaged in the first 600 nm close to the coverslip with a standard labeling of tubulin to visualize their microtubule network (Fig. 3a-3c). As evidenced in the xz cross section of 2 individual microtubules (Fig. 3d), the precision of ModLoc reveals the cylindrical structure of the microtubules. Although visualizing hollownesses is impeded by the labelling density and the spatial fluctuations of the microtubule directions (Fig. 3e), statistically significant sampling of several microtubule sections at various depths still resolves the cylindrical structure (Fig. 3f). In addition, its associated radial molecular density exhibits a central dip with a radius of ~12.3 nm as expected from the microtubule structure (Fig. 3g, see Supplementary Fig. 6). The tail at higher radii originates from the distance uncertainty introduced by the labelling.

**Figure 3.**
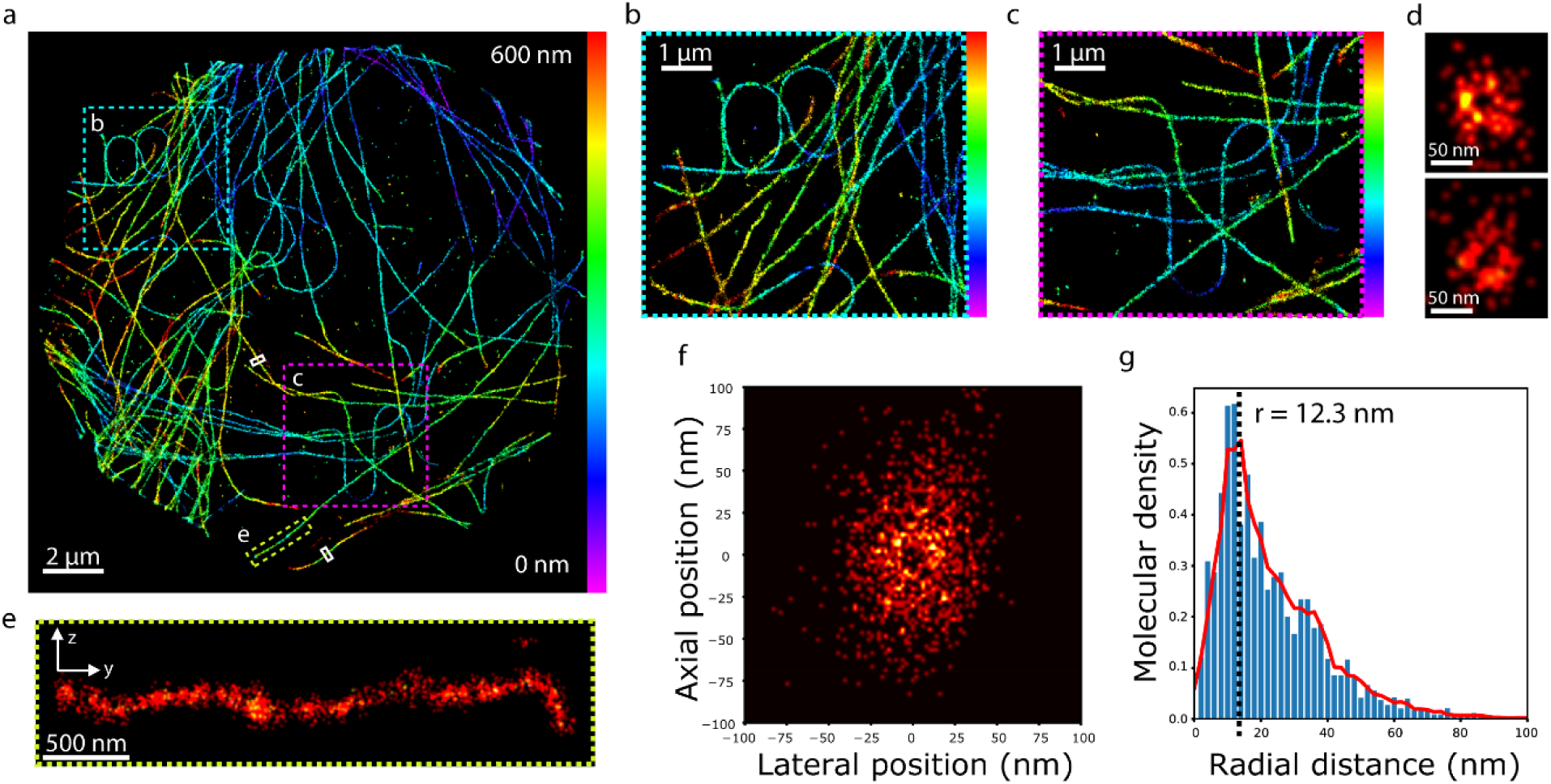
3D imaging of microtubules network in the first 600 nm in COS7 cell. **a-biological samples with ModLoc.** Full 3D imaging of tubulin in cos7 cell at the coverslip labeled with AF 647 fluorescent probes, **b** zoom on the cyan ROI of **a**. **c** zoom on the pink ROI in **a**. **d** Different yz cross sections highlighted by white ROI in **a**, which evidenced the cylindrical structure microtubules. **e** zoom on the yellow ROI of **a**, yz cross section of a microtubule**. f** Density map of detected fluorescent molecules for 7 different microtubules located at various depths within the capture range. **g** radial molecular density of **f** exhibiting a central dip with a radius of ~12.3 nm.

ModLoc is expected to achieve high performances in depth (Fig. 2f). Here we image various biological structures at depths of several microns. Figure 4a shows clathrin pits repartition in COS7 cell labeled with AF 647 at 4 microns in depth revealing the presence of the underlying nucleus. Cross sectional yz view of clathrin pits at various depth (Fig. 4b) show similar morphologies as the one obtained near the coverslip (Fig. 4c) confirming the constant axial localization precision of ModLoc. The performances of ModLoc for deep imaging have been further applied to mitochondria at 6 microns in depth in COS7 cell (Fig. 4d-4e). The measured cross sections show no difference in thickness (Fig. 4e) for the lower and the upper membranes of the mitochondria confirming the ModLoc constant axial precision over the whole capture range.

**Figure 4.**
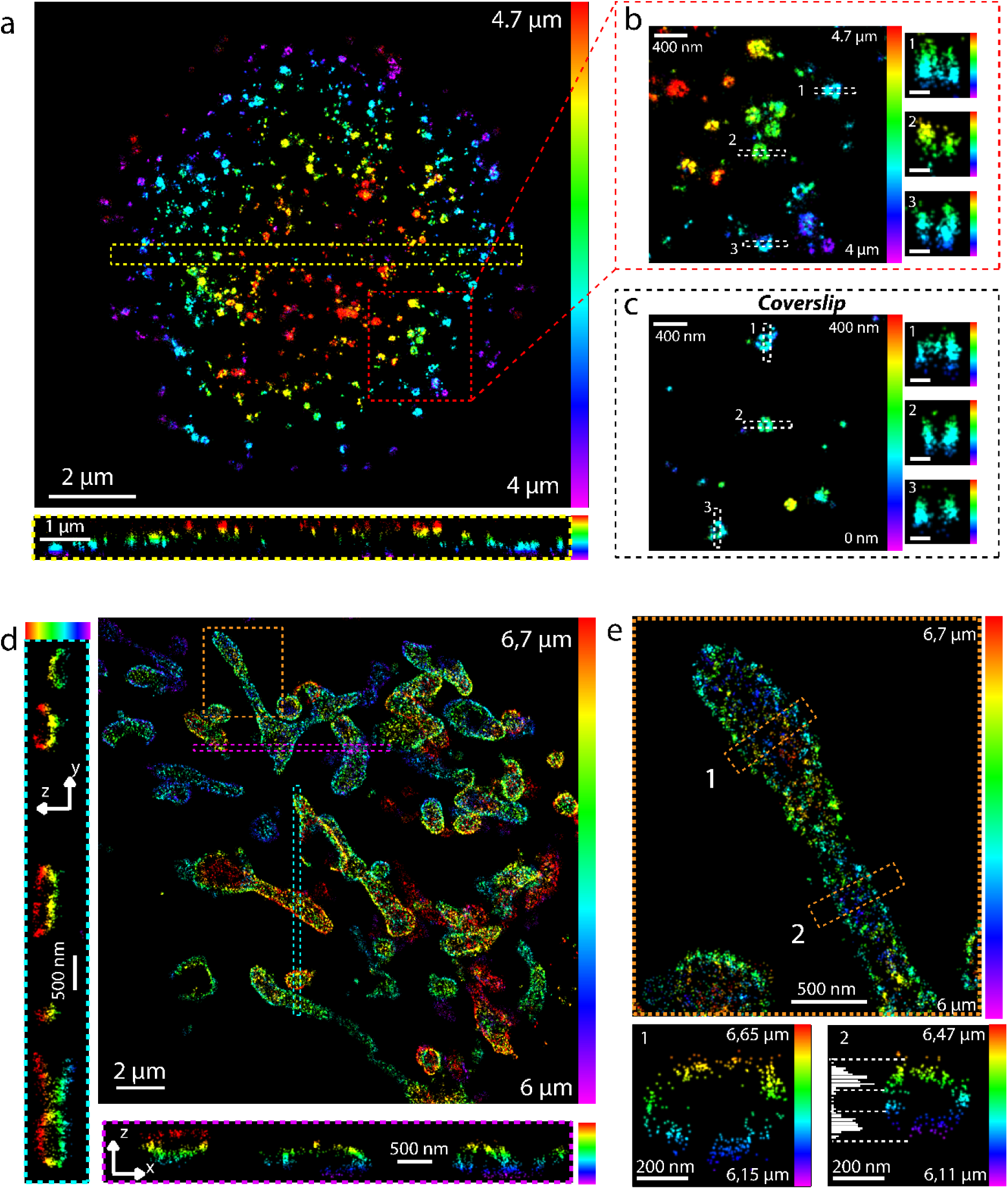
3D imaging of biological samples with ModLoc. **a**, <u>Top:</u> full 3D imaging of clathrin in cos7 cell at 4 microns in depth with heavy chain labeled with AF 647 fluorescent probes,. <u>Bottom:</u> xz cross section of the yellow ROI. The z bar color is defined in the same interval as in the top. **b**, <u>Left:</u> zoom on the red ROI of **a**. <u>Right:</u> different xz and yz cross sections of the white ROI. **c**, <u>Left:</u> 3D image of clathrin in cos7 at the coverslip with heavy chain labeled with AF 647 fluorescent probes. <u>Right:</u> different xz and yz cross sections of the white ROI. **d**, 3D image of mitochondria in cos7 cell at 6 microns in depth. Tom22 was labeled with AF 647 and 2 cross sections of ROIs are shown at left and at the bottom of the full field image. The colorbar range is the same for the 3 images. **e**, <u>Top:</u> zoom on the orange ROI of **d**. <u>Bottom 1 and</u> 2 correspond to xz cross sections at different positions along one mitochondria, with axial histogram showing a uniform z precision comparing the weighted standard deviation of the top and the bottom axial histogram distributions (respectively 30 nm and at the bottom is 32 nm).

We now evaluate the performances of ModLoc when facing harsh conditions at even higher depths and in the presence of strong optical aberrations. To perform observations at different depths, we use COS7 cells cultured in a collagen matrix with standard labeling of tubulin to visualize their microtubule network (see schematics in Fig. 5a). The wide field image of the microtubule network located 30 µm deep (taking into account the focal shift^37^) clearly shows the incidence of the aberrations and scattering induced by the collagen matrix (Fig. 5b). The axial information can still be retrieved with accuracy at such large depths as evidenced by the 3D reconstructed super-resolved image in Fig. 5c and close-ups in Fig. 5d and 5e. An axial projection of 3 microtubules on top of each other (Fig. 5f) shows that they are easily distinguishable when set 200 nm apart. Figure 5g presents the fluorophore localization in the cross section of 10 different microtubules at different (color coded) depths within the capture range. The localizations are homogeneously distributed along the lateral and axial direction confirming the constant performance of ModLoc within the capture range. The hollowness cannot be observed in such harsh conditions but the lateral and axial distributions have a gaussian shape related to the radial fluorophore profile around the microtubule (Fig. 5g, see Supplementary Figure 8). Measurements gives a lateral localization precision of about 15 nm and an axial precision of around 35 nm. The strong scattering properties of the collagen matrix impacts the localization precision in all directions.

**Figure 5.**
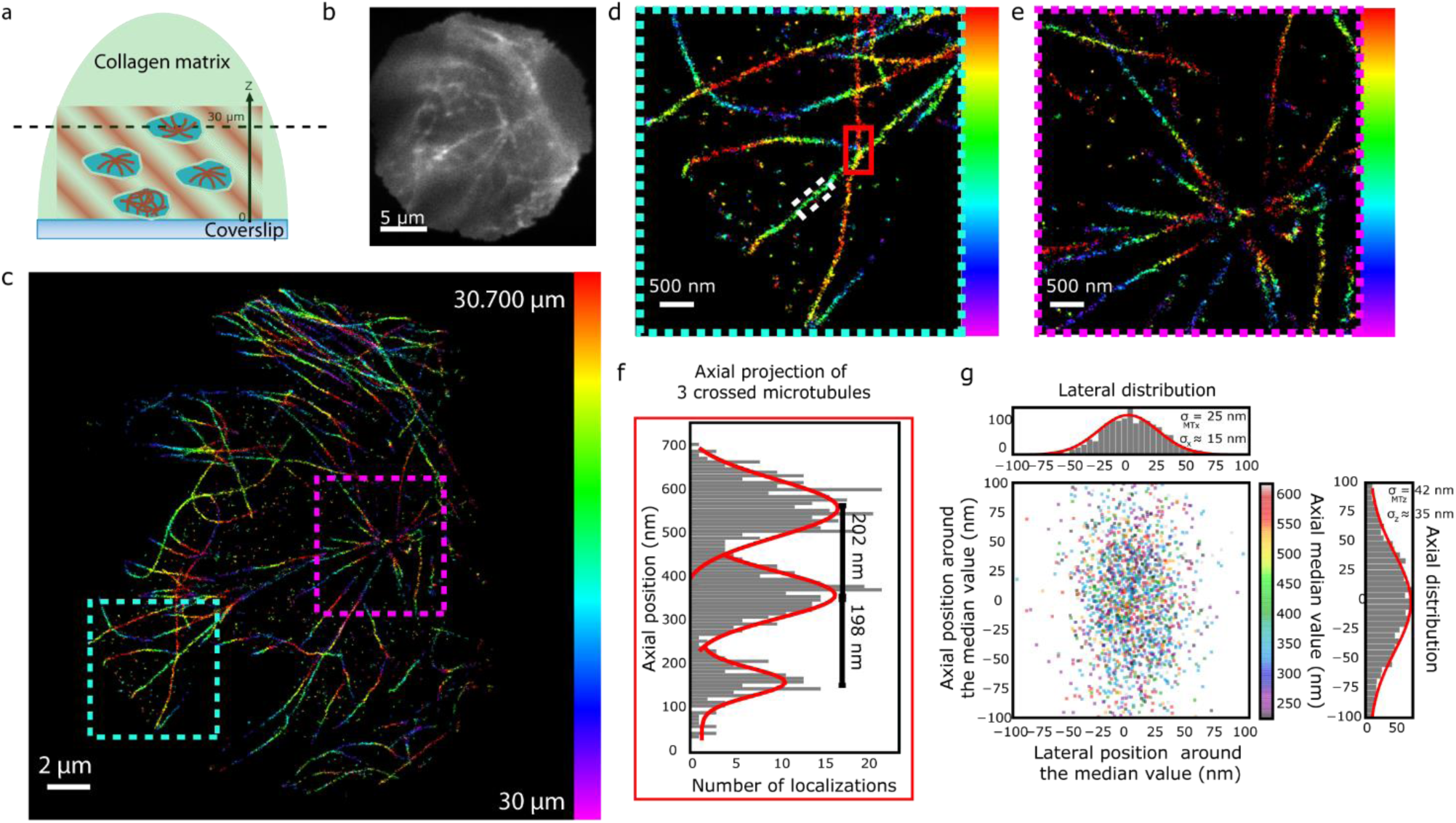
3D imaging of COS 7 cell microtubule grown in collagen matrix. **a**, Schematic representation of COS 7 grown in a collagen matrix. **b**, Wide field image of COS 7 cell tubulin at 30 microns in depth. **c**, 3D super resolved imaged of wide field represented in **b**. **d**, Zoom on the blue ROI in c. e, Zoom on the purple ROI in c. f, Axial cross section of 3 microtubules (red ROI in c) at different depths in the capture range. g, Scatter plot represents projection of different microtubules (supplementary for ROI positions) along around 500 nm for each. The median axial position of each microtubule in encoded with the color bar and lateral and axial distribution are represented with histograms. Weighted standard deviation of each histogram σ_MTx_ ~25 nm and σ_MTz_~42 nm gives an approximation of ModLoc localization precision σ_x_ ~15 nm and *σ*_z_~35 nm (Supplementary Figure 6).

We have demonstrated that ModLoc microscopy based on the introduction of a lock-in detection combining a tilted modulated excitation and a fast demodulation configuration offers real assets to full field 3D SMLM. This technique is easy to implement with a single objective configuration and offer a uniform axial localization precision at various depths. ModLoc technique takes full advantage of the robustness of the excitation pattern within the whole capture range as compared with PSF based techniques. ModLoc is well suited for deep biological samples imaging as demonstrated with cytoskeleton and various organelles images up to 30 µm.

ModLoc implementation could still be improved by modifying the excitation or the detection. The fringes pattern period could be reduced by using two counter propagating beams, either by using a reflecting mirror or in a 4Pi geometry which has the further advantage of doubling the collection efficiency^22^. In this case, the axial and lateral localizations become independent and an ultimate axial localization precision of *σ*_*ModLoc*_ ≈ 0.5 *σ*_*x*_ (see eq. 1 with *α*/2 = *θ* = 90°). The demodulation detection could use moving optical elements like mirrors which would be polarization independent and enhance the contrast albeit limited in frequency and potential source of mechanical drifts.

It is interesting to compare ModLoc with iPALM based approach. iPALM is based on the self-interference of the fluorescence while ModLoc uses the point-like nature of the fluorophore to report the phase of the illumination interference. The iPALM fundamental limit is thus determined by the same principles as ModLoc, the shot noise of the molecular fluorescence signal and the spatial period of the modulation. Phase wrapping and distortion of the PSF when imaging deeper in the sample restricted iPALM to thin samples and observations close to the coverslip, extension to deeper imaging was made possible only using adaptive optics^25^. Modloc has the advantage of being less sensitive to aberrations since its precision is not based on the propagation of a spherical wave but on two interfering planes waves producing its illumination pattern. Localization by modulating the excitation have been used in the early days of single emitter tracking along the lateral direction to probe myosin V stepping mechanism^38^ with gold nanobeads^39^. Recently, this strategy has been adapted to single molecule transverse localization for SMLM by various groups^40–42^. Similarly, MINFLUX^43^ technique is based on a structured illumination to localize the molecule but the estimator and the illumination pattern in the shape of a doughnut have been optimized to reach unmatched precision for a specific point (albeit breaking the translational invariance)^44^.

The next step would be to combine localization using illumination modulation in the three directions to retrieve 3D super-resolution imaging based only on the phase measurements. ModLoc phase detection is a generic concept used here to retrieve spatial information but it could also be extended to acquire additional information at the single molecule level like the molecular orientation or the local chemical environment through the fluorescence lifetime measurement.

**Methods are available in supplementary materials**

## Data availability

Data are available upon reasonable request to the corresponding author.

## Code avaibility

Processing code are based on already published solution as described in the supplementary

## Acknowledgements

P.J. acknowledge a master funding form GDR ImaBio, and PhD funding from IDEX Paris Saclay (ANR-11-IDEX-0003-02). M.B. is funded by the Labex PALM. We acknowledge the contribution of the Centre de Photonique BioMédicale to cell culture and labeling. We also thank Guillaume Dupuis for discussion and Surabhi Sreenivas for a careful reading of the paper. We thank Abbelight for the free use of NEMO software and dSTORM buffers. This work was supported by the AXA research fund, the ANR (LABEX WIFI, ANR-10-LABX-24), ANR MSM-Modulated super-resolution microscopy (ANR-17-CE09-0040), the valorization program of the IDEX Paris Saclay and of Labex PALM.

## Author contributions

P.J, C.C., N.B., C.P., E.F. and S.L.F. conceived the project. P.J. designed the optical setup, performed the acquisitions, CRLB calculations and data analysis. P.J. and E.F. carried out simulations. N.B. developed the (d)STORM buffer. N.B., C.C. and P.J. optimized the immunofluorescence protocol. P.J., C.C. and S.L.F prepared the COS-7 cells samples. M.B designed the 3D sample protocol. All authors contributed to writing the manuscript.

