## supplementary material for "Nanometric axial localization of single fluorescent molecules with modulated excitation"

**Supplementary Figure 1 :** Optical setup of ModLoc.

**Supplementary Figure 2 :** Schematic of the temporal intensity distribution on the four sub-arrays of the camera.

**Supplementary Figure 3 :** Schematic of the pattern orientation and the z information encoded.

**Supplementary Figure 4 :** Cramér-Rao Lower Bounds for different modulation contrasts.

**Supplementary Figure 5 :** Calibration of the fringe pattern.

**Supplementary Figure 6 :** Regions of interest of the different microtubule profiles at the coverslip.

**Supplementary Figure 7 :** Regions of interest of the different microtubule profiles at 30 microns.

**Supplementary Figure 8 :** Localization precision estimation from tubulin profiles.

**Supplementary Note 1 :** Optical setup description

**Supplementary Note 2 :** Fluorescence transmission in the four sub-arrays.

**Supplementary Note 3 :** Cramér-Rao Lower Bounds.

**Supplementary Note 4 :** Excitation pattern calibration.

**Supplementary Note 5 :** Localization precision measurement and calibration samples

**Supplementary Note 6 :** Sample preparation

**Supplementary Note 7 :** Data workflow processing

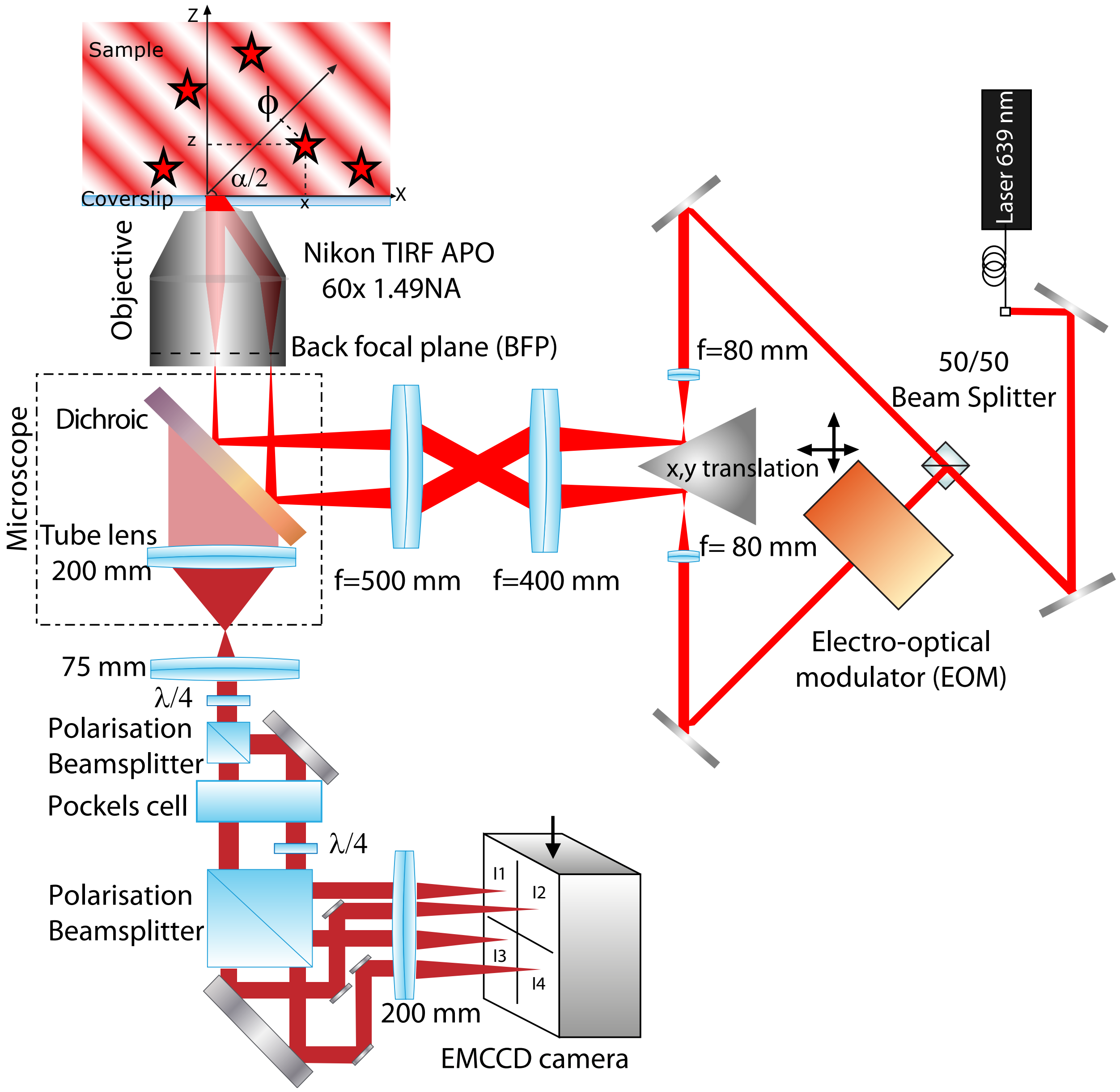


**Supplementary Figure 1: Optical setup of ModLoc**. More details in **Supplementary Note 1**

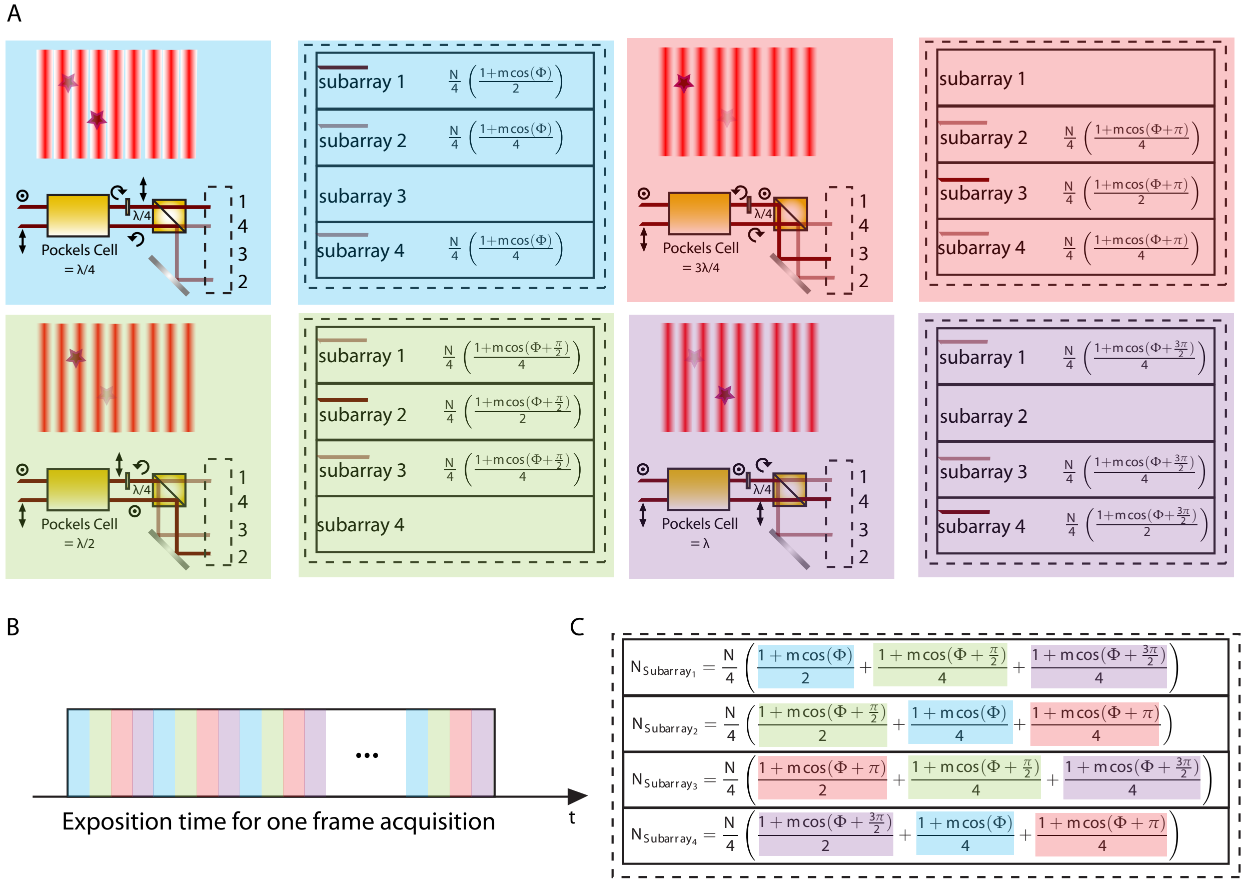


**Supplementary Figure 2: Schematic of the temporal intensity distribution on the four sub-arrays of the camera. (A)** Intensity distribution for each Pockels cell state. N is the photon number emitted by a single emitter. **(B)** Chronogram of the Pockels cell states during one acquisition frame. **(C)** Final intensity for each sub-array of the camera at the end of the acquisition. See **Supplementary Note** 2.

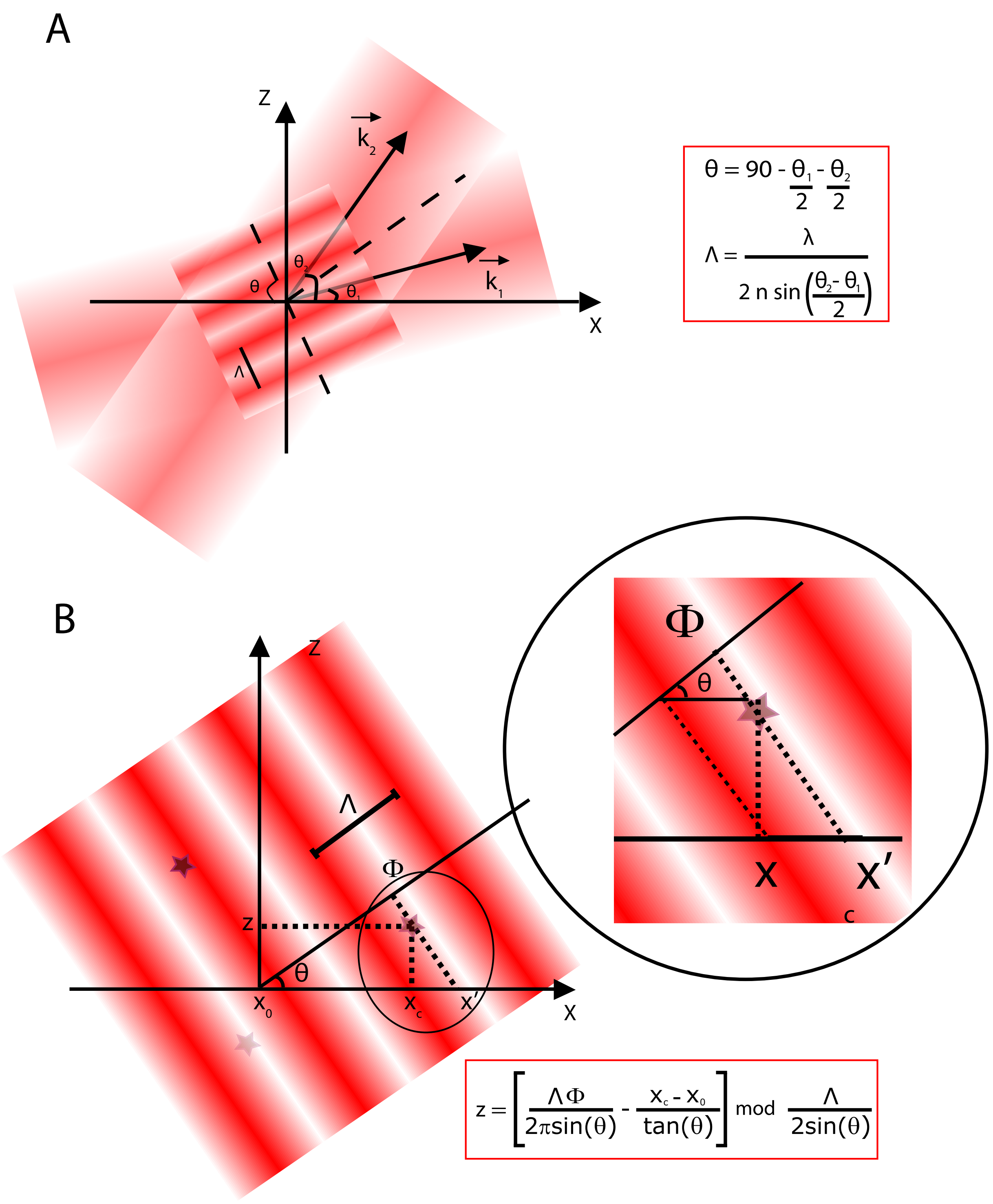


**Supplementary Figure 3: Schematic of the pattern orientation and the z information encoded. (A)** Schematic representation of the fringe pattern orientation as a function of the 2 incident angles of the 2 interfered waves. The z tilt of the fringe pattern is given by the simple equation $\theta = \frac{\pi}{2} - \frac{\theta_2}{2} - \frac{\theta_1}{2}$. The spatial frequency of the fringe pattern is given by $\Lambda = \frac{\lambda}{2n\sin(\frac{\theta_2-\theta_1}{2})}$, where $\lambda$ is the wavelength of the excitation, $n$ is the medium index and $\theta_1$ and $\theta_2$ the angles of the 2 interfering waves. **(B)** Representation of the information extract from a single probe. The transverse position is obtained from classical gaussian fitting and the z information is obtained from the combinaison of the transverse position and the phase information. The z projection of the spatial frequency must be about the depth imaging in order to avoid unwrapping uncertainty.

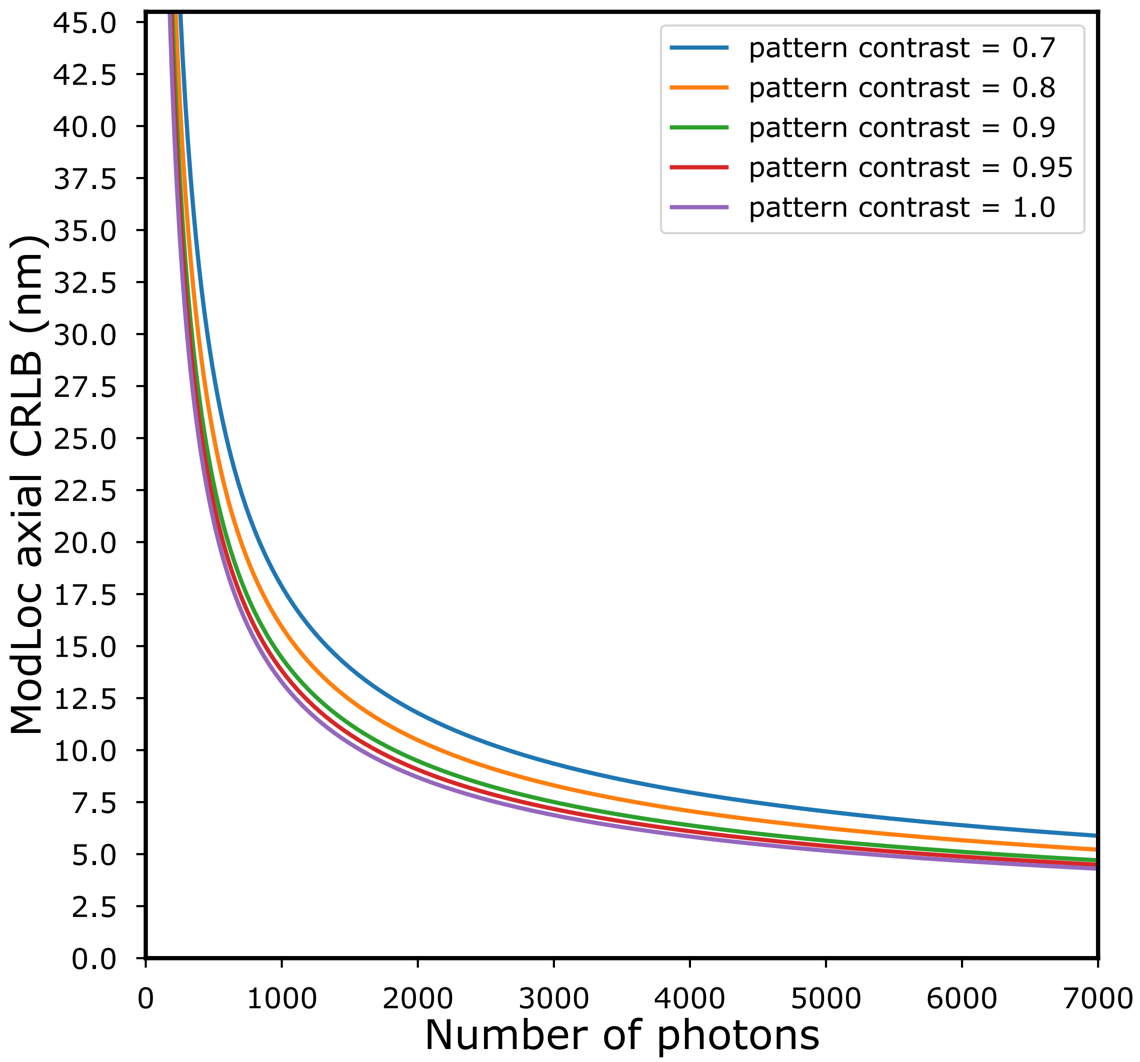


**Supplementary Figure 4: Cramér-Rao Lower Bounds of ModLoc for optimum pattern parameters as a function of the number of photons.** Cramér-Rao Lower Bounds variations with the number of photons by taking a fringe angle is about 49° and a spatial frequency of the fringe pattern about 534 nm in order to get the axial fringe projection to 700 nm. See **Supplementary Note 3** for more details.

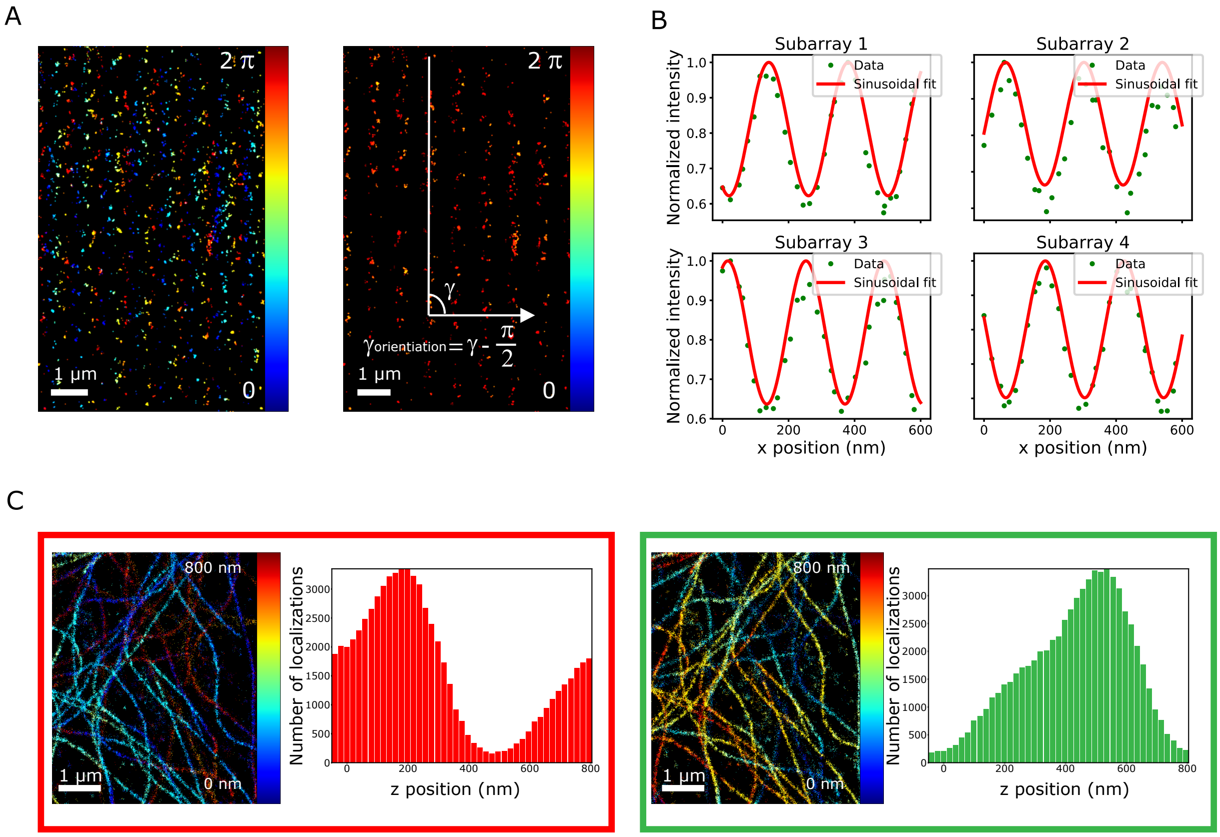


**Supplementary Figure 5: Techniques used for the pattern calibration . (A)** Phase images used to obtain the angle orientation of the fringe pattern in the xy plane. The iso-phase selection correlate with a model fringe pattern gives access to the orientation angle. **(B)** Raw signal and sinusoidal model fitting of a fluorescent probe intensity. The sample displacement along the modulation direction modulates the fluorescence signal of the 40-nm-diameter red nanobead. The spatial frequency of the fitted signal correspond to the the spatial frequency of the fringe pattern. The signal is represented for the 4 sub-arrays, the phase shift between the reconstructed signal corresponds to the 4 fringe positions. **(C)** Reconstructed images and their z distribution corresponding of the same region of interest for 2 different pattern origins. The red part corresponds to a 3D reconstructed image with an inaccurate pattern origin. Note that the z distribution is wrapped. The green part corresponds to the same area, reconstructed with the right pattern origin obtained with the z distribution information. See **Supplementary Note** 4 for more details.

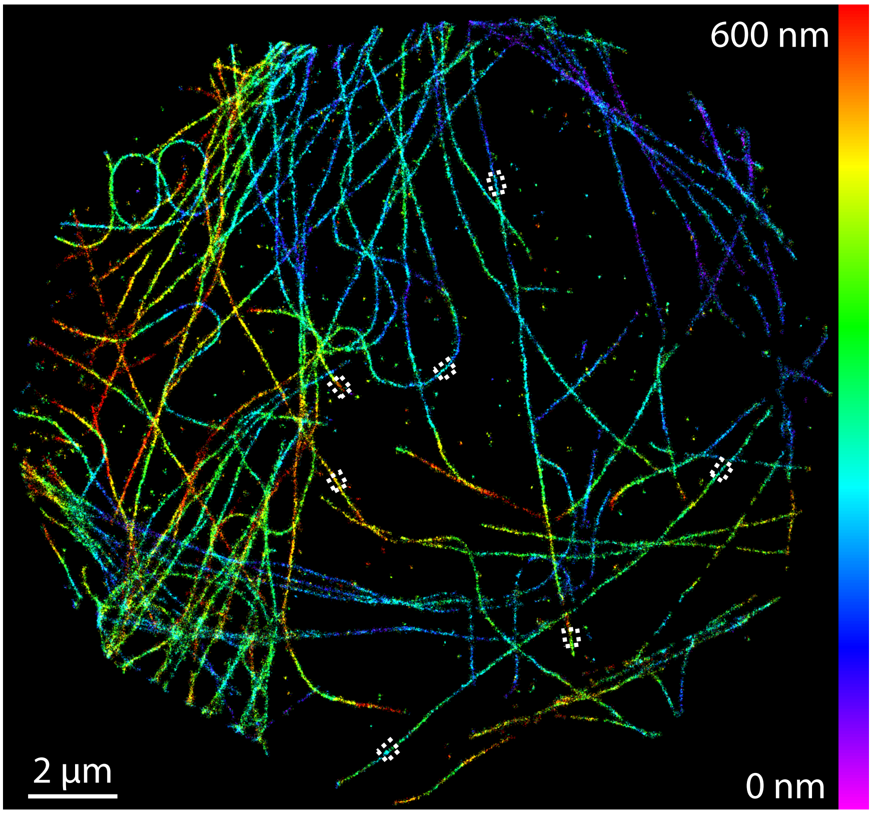


**Supplementary Figure 6:** Regions of interest (ROIs) of the different microtubule profiles presented in the figure 3 (coverslip). Integration of all ROIs had been performed over 200 nm

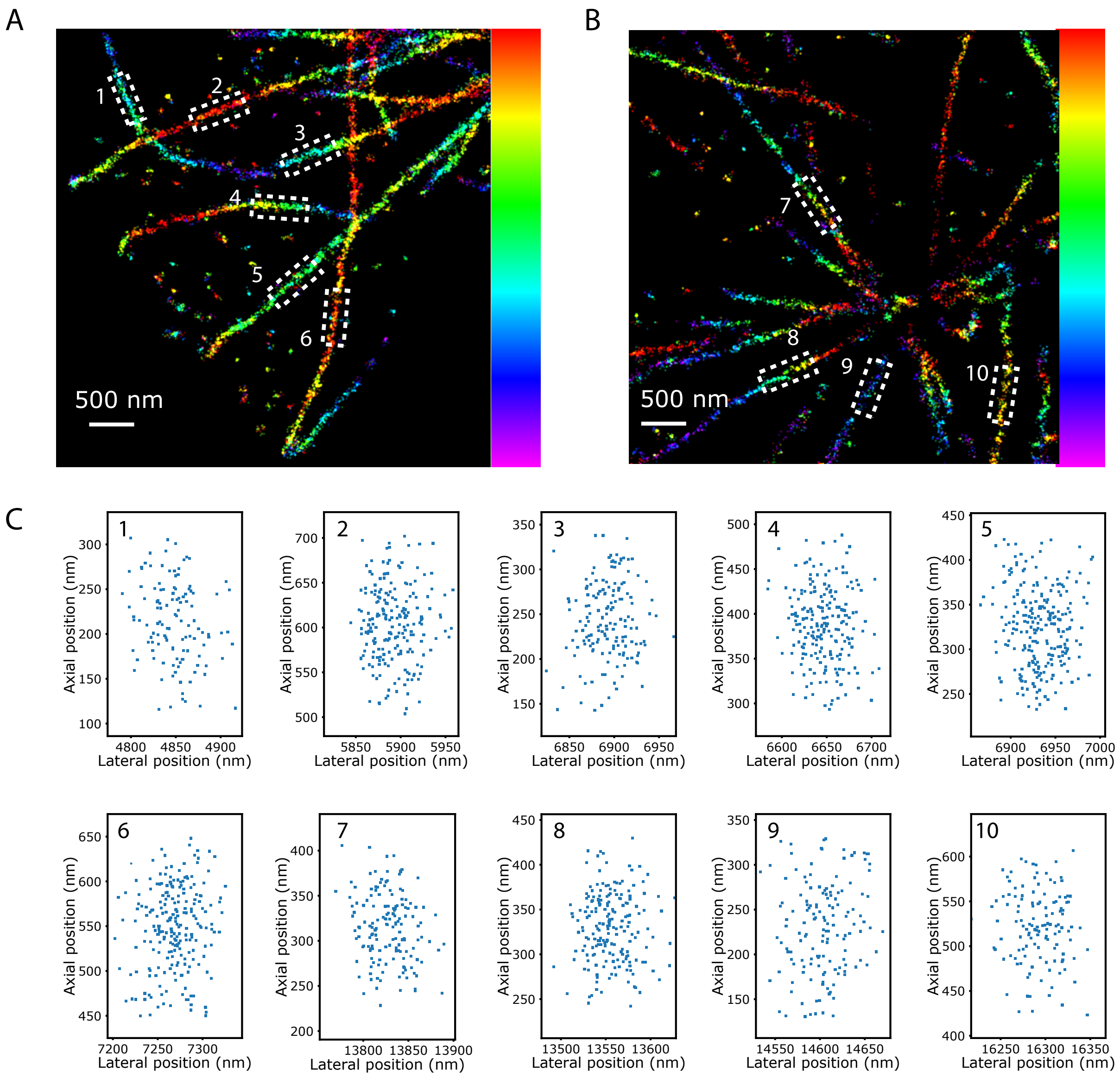


**Supplementary Figure 7:** Regions of interest of the different microtubule profiles presented in the figure 4 (30 microns). **(A)** Zoom on the first area. The different ROI are represented in white. Integration of all ROIs had been performed over 500 nm. **(B)** Zoom on the second area. The different ROI are represented in white. Integration of all ROIs had been performed over 500 nm. **(C)** Individual cross sections of the 10 selected ROIs.

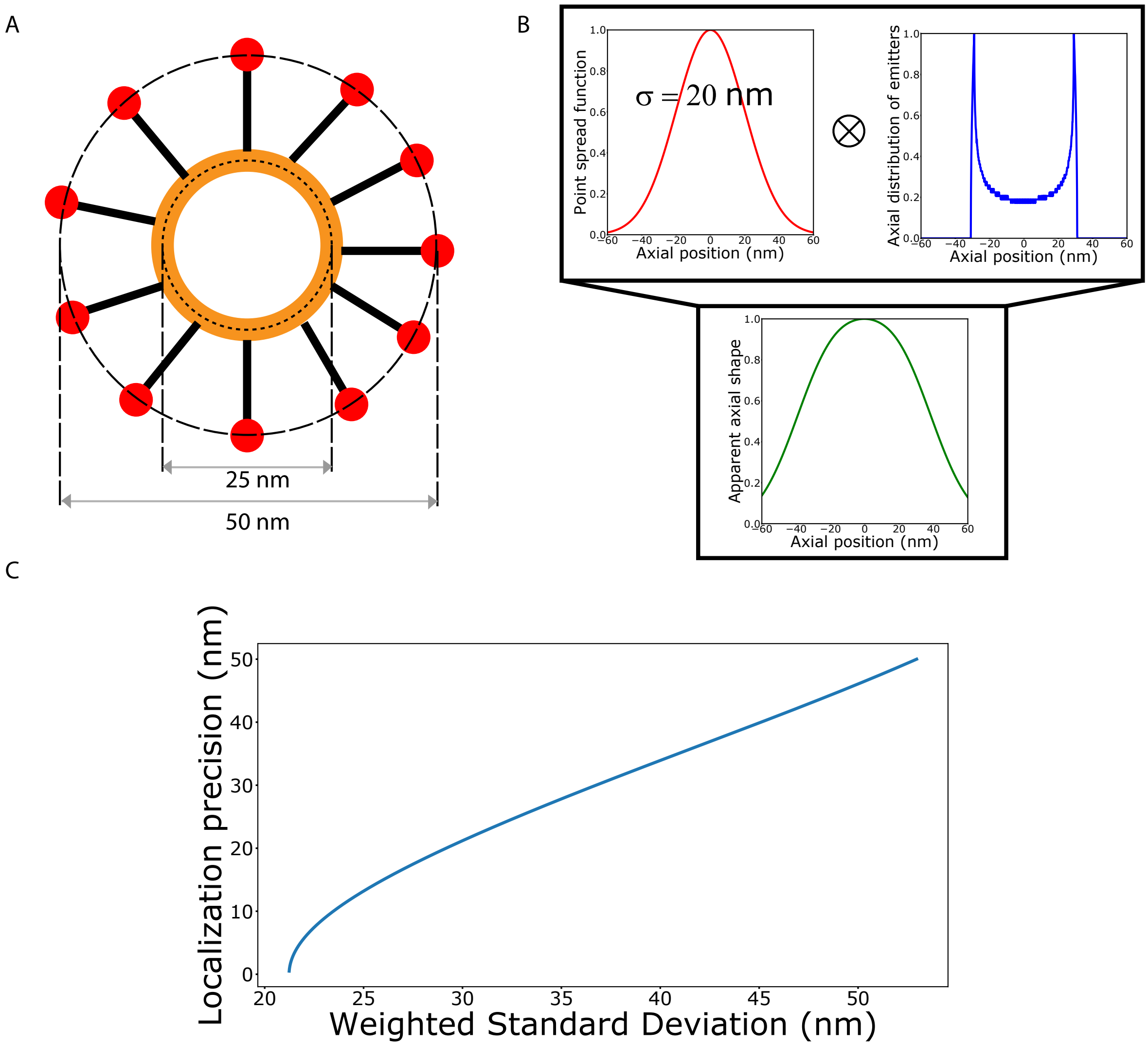


**Supplementary Figure 8:** Localization precision estimation from tubulin profiles. **(A)** Schematic of a microtubule (orange disks) immunolabeled with a primary antibody and a fragment (Fab2) of secondary antibody labeled with Alexa 647 (red disks). Localization precision of ModLoc had been experimentally determined on microtubules at 30 microns. The apparent shape of a microtubule does not only depend on the localization precision but also on the size of linkers used for immunolabelling [1]. **(B)** Convolution of the theoretical transverse distribution of fluorescent probes with a Gaussian function corresponding to the PSF of an emitter. By considering the size of the biological structure and the size of the labeling, the effective size of a microtubule is 60 nm. The PSF size is related to the localization precision of the method used , the apparent shape of a microtubule can be approximated by convoluting the PSF of an emitter with the spatial distribution along one axis. **C** Localization precision as a function of the weighted standard deviation of the resulting distribution.

### Supplementary Note 1 : Optical setup

A complete schematic of the optical setup used is presented in **Supplementary Figure** 1 . We used a Nikon Eclipse Ti inverted microscope with a Nikon Perfect Focus System. The dichroic and filters in the microscope cube are from Semrock (Di03-R635-t1-25x36 and BLP01-635R-25). The excitation was performed using a 639-nm laser (Genesis MX 639, 1W, Coherent), couple to a single-mode optical fiber maintaining polarization. The laser beam was separated in 2 paths and recombined in the sample through a Nikon APO TIRF x60 1.49 NA oil immersion objective lens. The fluorescence was collected through the same objective lens, sent in the demodulation module and recorded on four quadrants of a 512x512-pixel EMCCD camera (iXon3, Andor). The camera was placed at the focal plane of the module of magnification 2.6 and the optical pixel size was approximately 100 nm. Finally, the imaging paths were calibrated in intensity to compensate the non-ideality of the transmission for all channels. The phase modulation is the excitation path is performed by an electro optical modulator (EOM) (EO-PM-NR-C1, Thorlabs), while the phase sorting in the detection path is performed by a Pockels cell (CF1043-20SG-500/700, Fast pulse) controlled by a high voltage power amplifier (10/10B-HS, TREK). The synchronization was provided thanks to a 4-channel generator that triggers the EOM, the Pockels cell and the camera.

### Supplementary Note 2 : Fluorescence transmission in the four sub-arrays

The camera used to collect the emitted fluorescence is divided in four sub-arrays in order to detect the four samples of the modulated emission signal in a single acquired image. The fluorescence first passes through a $\lambda/4$ waveplate and is divided in 2 paths thanks to a polarizing beam splitter. Because the fluorescence emission of the fluorophore is not polarized with the currently used dyes' linker, the two paths have the same intensity but two orthogonal linear polarizations. The demodulation module also comprises a Pockels cell common to these two paths, with specific axes oriented at 45 °regarding the two orthogonal polarization paths. After the Pockels cell, another $\lambda/4$ waveplate is inserted in only one of the paths. One of the polarizations thus becomes circular while the other stays linear for all Pockels cell states. The last polarizing component is a polarizing beam splitter that separates the initial signal in four paths. These four paths are collected on the camera in different sub-arrays, which implies that each sub-array receives no signal for one fringe position.

For the first fringe position, the Pockels cell acts as a $\lambda/4$ plate and half of the photons are detected in sub-array 1 while sub-arrays 2 and 4 each receive a quarter of the signal. For the second fringe position, the Pockels cell acts as a $\lambda/2$ plate and half of the photons are detected in sub-array 2 while sub-arrays 1 and 3 each receive a quarter of the signal. Similarly, when the Pockels cell corresponds to a 3 $\lambda/4$ plate, half of the photons are detected in sub-array 3 while sub-arrays 2 and 4 each receive a quarter of the total signal. Finally, sub-array 4 receives half of the photons while sub-arrays 1 and 3 each receive a quarter corresponding to a pockels cell state of $\lambda$. Assuming N photons emitted by a fluorescent probe, the total signal on each channel reads :

$$I_{Subarray_1} = \frac{N}{4}\left(\frac{1+m\cos(\Phi)}{2} + \frac{1+m\cos(\Phi+\frac{\pi}{2})}{4} + \frac{1+m\cos(\Phi+\frac{3\pi}{2})}{4}\right) \tag{S.1a}$$

$$= \frac{N}{4}\left(\frac{1+m\cos(\Phi)}{2} + \frac{1}{2}\right) \tag{S.1b}$$

$$I_{Subarray_2} = \frac{N}{4}\left(\frac{1+m\cos(\Phi+\frac{\pi}{2})}{2} + \frac{1+m\cos(\Phi)}{4} + \frac{1+m\cos(\Phi+\pi)}{4}\right) \tag{S.2a}$$

$$= \frac{N}{4}\left(\frac{1+m\cos(\Phi+\frac{\pi}{2})}{2} + \frac{1}{2}\right) \tag{S.2b}$$

$$I_{Subarray_3} = \frac{N}{4}\left(\frac{1+m\cos(\Phi+\pi)}{2} + \frac{1+m\cos(\Phi+\frac{\pi}{2})}{4} + \frac{1+m\cos(\Phi+\frac{3\pi}{2})}{4}\right) \tag{S.3a}$$

$$= \frac{N}{4}\left(\frac{1+m\cos(\Phi+\pi)}{2} + \frac{1}{2}\right) \tag{S.3b}$$

$$I_{Subarray_4} = \frac{N}{4}\left(\frac{1+m\cos(\Phi+\frac{3\pi}{2})}{2}+\frac{1+m\cos(\Phi)}{4}+\frac{1+m\cos(\Phi+\pi)}{4}\right) \tag{S.4a}$$

$$= \frac{N}{4}\left(\frac{1+m\cos(\Phi+\frac{3\pi}{2})}{2}+\frac{1}{2}\right) \tag{S.4b}$$

59 A schematic of the modulation is represented in **Supplementary Figure** 2

### Supplementary Note 3 : Fisher information and Cramér-Rao Lower Bounds

The Cramér-Rao lower bounds (CRLB) is the most commonly used tool to obtain the theoretical localization precision achievable by a technique. It is obtained from the Fisher Matrix, constructed from the likelihood function of our information $\vec{n}$ : the number of photons detected from our camera.

$$I_{u,v}(\Theta) = E[\frac{\partial \ln L(\vec{n}|\Theta)}{\partial \Theta_u} \frac{\partial L(\vec{n}|\Theta)}{\partial \Theta_v}] \tag{S.5}$$

The z information is obtained from the ModLoc information based on the single emitter position inside the fringe pattern, and its lateral position obtained from Gaussian fitting. It is possible to obtain the CRLB information for the z position by considering the uncertainty of both techniques. The z position is obtained from the formula :

$$z = \frac{\Lambda \phi}{2\pi \sin(\theta)} - \frac{x - x_0}{\tan(\theta)} \tag{S.6}$$

Where $\theta$ is the angle of the tilted fringe pattern with the lateral plane, $x_0$ is the origin of the fringe fringe pattern, $x$ the lateral position obtained from Gaussian fitting and $\phi$ the phase of the modulated signal from ModLoc method, defined as $\phi = \frac{2\pi}{\Lambda} r$, $r$ is the position along the fringe direction. From this equation, it is obvious that the Cramér-Rao lower bounds can be obtained as follow :

$$\sigma_z^{CRLB} = \sqrt{\left(\frac{\partial z}{\partial r}\right)^2 \sigma^2_{r,CRLB} + \left(\frac{\partial z}{\partial x_c}\right)^2 \sigma^2_{x_c,CRLB}} \tag{S.7a}$$

$$\sigma_z^{CRLB} = \sqrt{\frac{\sigma^2_{r,CRLB}}{\sin^2(\theta)} + \frac{\sigma^2_{x_c,CRLB}}{\tan^2(\theta)}} \tag{S.7b}$$

The phase and position error then define the minimum theoretical localization precision that we can obtain with ModLoc. These error are calculated in the following development.

#### 1 Experimental model of a detected Point Spread Function

To begin, it is essential to consider the theoretical model of our information. We can assimilate the Point Spread Function (PSF) to a simplified form like an isotropic Gaussian Function as follow :

$$PSF(x,y) = \frac{N}{2\pi\sigma^2} \exp - \frac{(x - x_c)^2 + (y - y_c)^2}{2\sigma^2} \tag{S.8}$$

Where $x_c$ and $y_c$ are the real position of the fluorescent emitter and $N$ is the total number of photons emitted. We use the same approach as [2] to take into account the PSF sampling on the camera. In this case, the PSF is sampled over an area composed of several pixel of side note $a$. We write the signal received by a pixel as :

$$\nu(x,y) = \int_{-\frac{a}{2}}^{\frac{a}{2}} \int_{-\frac{a}{2}}^{\frac{a}{2}} PSF(x,y)dxdy \tag{S.9}$$

We note that $\int_0^x \exp -t^2 dt = \frac{\sqrt{\pi}}{2}\operatorname{erf}(x)$. Doing a simple change of variables, equation (S.9) can be expressed as :

$$\nu(x,y) = N\Delta E(x)\Delta E(y) \tag{S.10}$$

where

$$\Delta E(u) = \frac{1}{2}[\operatorname{erf}\left(\frac{u-u_c+\frac{a}{2}}{\sqrt{2}\sigma}\right) - \operatorname{erf}\left(\frac{u-u_c-\frac{a}{2}}{\sqrt{2}\sigma}\right)] \tag{S.11}$$

This expression corresponds to a perfect case without noise background impact. In order to facilitate the analysis, we consider a uniform Background noise $b$. The pixel intensity model simply becomes :

$$\nu(x,y) = N\Delta E(x)\Delta E(y) + b \tag{S.12}$$

The difference between our model and the classical model comes from the fact that the number of photon emitted by the emitter depends of its position in the structured pattern. The photon emission is temporally modulated with a fringe pattern applied along the $\vec{r}$ direction. Our goal is to demodulate this signal with several samples. If we consider 4 samples corresponding to 4 fringe positions, the expression of the pixel $j$ at the fringe position $i$ ( $i = 0, 2...3$) is given by :

$$\nu_j^i = N^i(r)\Delta E(x_j)\Delta E(y_j) + \frac{b}{4} \tag{S.13}$$

with the different values of $N^i(x_c)$ :

$$N^i(r) = \frac{N}{8}\left(2 + m\cos(\frac{2\pi}{\Lambda}r + i\frac{\pi}{2})\right) \tag{S.14}$$

where $m$ is the pattern contrast, $\Lambda$ is the spatial frequency of the fringe pattern and $r$ is the position along the fringe direction. Note that we consider only noise coming from sample, the camera noise is neglected (the EMCCD read out noise is less than 1e rms, the offset and gain per pixel are not considered too).

#### 2 Likelihood function and Cramér-Rao Lower Bounds of ModLoc

Knowledge of the likelihood function is required in order to construct the fisher matrix. Considering a measurement of photon $n$ on the pixel $j$. The collected data follows a poisson law as :

$$p(n^i_j) = \frac{\nu^i_j(r,x,y)^{n^i_j}\exp(-\nu^i_j(r,x,y))}{n^i_j!} \tag{S.15}$$

The number of photons $N^i$ collected from the PSF corresponding to the fringe position i is the sum of collected data $\vec{n^i} = [n^i_0, n^i_1, ....n^i_j]$ over $M$ pixels composing the ROI : $N^i = \sum_{j=0}^{M} n^i_j$. We call $\Theta$ the parameter vector composed of $\Theta = [r, x_c, y_c, N, b]$. Therefore, we can write the likelihood function $L(\vec{n^i}|\Theta)$ and the log-likelihood function $l(\vec{n^i}|\Theta) = \ln L(\vec{n^i}|\Theta)$ of our model as :

$$L(\vec{n^i}|\Theta) = \prod_{i=0}^{3}\prod_{j=0}^{M} p(n^i_j) \tag{S.16a}$$

$$L(\vec{n^i}|\Theta) = \prod_{i=0}^{3}\prod_{j=0}^{M} \frac{\nu^i_j(r,x,y)^{n^i_j}\exp(-\nu^i_j(r,x,y))}{n^i_j!} \tag{S.16b}$$

$$l(\vec{n^i}|\Theta) = \sum_{i=0}^{3}\sum_{j=0}^{M} n^i_j \ln(\nu^i_j(r,x,y)) - \nu^i_j(r,x,y) - \ln n^i_j! \tag{S.16c}$$

The Fisher matrix is obtained from partial derivatives of the log-likelihood function, define from the likelihood function as $l(\vec{n^i}|\Theta) = \ln L(\vec{n^i}|\Theta)$. We can then rewrite equation (S.5) :

$$I_{u,v}(\Theta) = E[\sum_{i=0}^{3}\sum_{j=0}^{M}\left(\frac{n^i_j - \nu^i_j(r,x,y)}{\nu^i_j(r,x,y)}\right)^2 \frac{\partial\nu^i_j(r,x,y)}{\Theta_u}\frac{\partial\nu^i_j(r,x,y)}{\Theta_v}] \tag{S.17a}$$

$$I_{u,v}(\Theta) = \sum_{i=0}^{3}\sum_{j=0}^{M} \frac{1}{v_j^i(r,x,y)} \frac{\partial v_j^i(r,x,y)}{\Theta_u} \frac{\partial v_j^i(r,x,y)}{\Theta_v} \tag{S.17b}$$

With partial derivatives :

$$\frac{\partial v_j^i(r,x,y)}{\partial r} = \Delta E(y)\Delta E(x)\frac{\partial N^i(r)}{\partial r} \tag{S.18a}$$

$$\frac{\partial v_j^i(r,x,y)}{\partial x_c} = N^i(r)\Delta E(y)\frac{\partial \Delta E(x)}{\partial x_c} \tag{S.18b}$$

$$\frac{\partial v_j^i(r,x,y)}{\partial y_c} = N^i(r)\Delta E(x)\frac{\partial \Delta E(y)}{\partial y_c} \tag{S.18c}$$

$$\frac{\partial v_j^i(r,x,y)}{\partial N} = \frac{1}{8}\left(2 + m\cos(\frac{2\pi}{\Lambda}r + i\frac{\pi}{2})\right)\Delta E(x)\Delta E(y) \tag{S.18d}$$

$$\frac{\partial v_j^i(r,x,y)}{\partial b} = \frac{1}{4} \tag{S.18e}$$

Both derivatives expression of $\frac{\partial N^i(r)}{\partial r}$ and $\frac{\partial \Delta E(u)}{\partial u_c}$ are developed :

$$\begin{aligned}
\frac{\partial \Delta E(u)}{\partial u_c} &= \frac{1}{2}\frac{\partial}{\partial u_c}\left(\mathrm{erf}\left(\frac{u-u_c+\frac{a}{2}}{\sqrt{2}\sigma}\right) - \mathrm{erf}\left(\frac{u-u_c-\frac{a}{2}}{\sqrt{2}\sigma}\right)\right)\\
\frac{\partial \Delta E(u)}{\partial u_c} &= \frac{1}{2}\frac{\partial}{\partial u_c}\left(\frac{2}{\sqrt{\pi}}\int_0^{\frac{u-u_c+\frac{a}{2}}{\sqrt{2}\sigma}} \exp(-u^2)du - \frac{2}{\sqrt{\pi}}\int_0^{\frac{u-u_c-\frac{a}{2}}{\sqrt{2}\sigma}} \exp(-u^2)du\right)\\
\frac{\partial \Delta E(u)}{\partial u_c} &= \frac{1}{\sqrt{\pi}}\frac{\partial}{\partial u_c}\left(\frac{1}{\sqrt{2}\sigma}\int_0^{u_c} \exp(-\left(\frac{u+\frac{a}{2}-t}{\sqrt{2}\sigma}\right)^2)dt - \frac{1}{\sqrt{2}\sigma}\int_0^{u_c} \exp(-\left(\frac{u-\frac{a}{2}-t}{\sqrt{2}\sigma}\right)^2)dt\right)\\
\frac{\partial \Delta E(u)}{\partial u_c} &= \frac{1}{\sqrt{2\pi}\sigma}\frac{\partial}{\partial u_c}\left(\int_0^{u_c} \exp(-\left(\frac{u+\frac{a}{2}-t}{\sqrt{2}\sigma}\right)^2)dt - \int_0^{u_c} \exp(-\left(\frac{u-\frac{a}{2}-t}{\sqrt{2}\sigma}\right)^2)dt\right)\\
\frac{\partial \Delta E(u)}{\partial u_c} &= \frac{1}{\sqrt{2\pi}\sigma}\left(\exp(-\left(\frac{u-u_c-\frac{a}{2}}{\sqrt{2}\sigma}\right)^2) - \exp(-\left(\frac{u-u_c+\frac{a}{2}}{\sqrt{2}\sigma}\right)^2)\right)
\end{aligned} \tag{S.19}$$

$$\begin{aligned}
\frac{\partial N^i(x_c)}{\partial r} &= \frac{\partial}{\partial r}\frac{N}{8}\left(2 + m\cos(\frac{2\pi}{\Lambda}r + i\frac{\pi}{2})\right)\\
\frac{\partial N^i(x_c)}{\partial r} &= -\frac{N}{4\Lambda}\pi m \sin(\frac{2\pi}{\Lambda}r + i\frac{\pi}{2})
\end{aligned} \tag{S.20}$$

These equations allow the final Fisher matrix expression. The main characteristic of the Fisher matrix is the obtaining of the variance lower bound of the considered parameter as $\mathrm{var}(\Theta_u) \geq \frac{1}{I_{uu}\Theta}$. By this way, it is possible to get the theoretical localization precision of our model : $\sigma_u^{CRLB} = \sqrt{\operatorname{var}\Theta_u}$

$$I_{rr}(\Theta) = \sum_{i=0}^{3}\sum_{j=0}^{M} \frac{1}{v_j^i(r,x,y)} \left(\frac{\partial v_j^i(r,x,y)}{\partial r}\right)^2 \tag{S.21a}$$

$$\sigma_r^{CRLB} = \sqrt{\left(\frac{1}{\sum_{i=0}^{3}\sum_{j=0}^{M} \frac{1}{v_j^i(r,x,y)} \left(\frac{\partial v_j^i(r,x,y)}{\partial r}\right)^2}\right)} \tag{S.21b}$$

Where $\sigma_r^{CRLB}$ is one part of the minimum theoretical localization precision used to obtained z Cramér-Rao lower bounds, and the previous expression are used to compute curve in (**Supplementary Figure** 4. In order to simply calculation, we will consider in the next development that the modulation contrast is equal to 1 and the background is equal to 0. By injecting (S.13), (S.18a) and (S.20) in (S.21a), we write :

$$I_{rr}(\Theta) = \sum_{j=0}^{M} \frac{(\Delta E(y)\Delta E(x))^2}{\Delta E(x_j)\Delta E(y_j)} \sum_{i=0}^{3} \frac{\left(-\frac{N}{4\Lambda}\pi \sin(\frac{2\pi}{\Lambda}r + i\frac{\pi}{2})\right)^2}{\frac{N}{8}\left(2+\cos(\frac{2\pi}{\Lambda}r + i\frac{\pi}{2})\right)} \tag{S.22a}$$

$$I_{rr}(\Theta) = \frac{N}{2\Lambda^2}\pi^2 \sum_{i=0}^{3} \frac{\sin^2(\frac{2\pi}{\Lambda}r + i\frac{\pi}{2})}{\left(2+\cos(\frac{2\pi}{\Lambda}r + i\frac{\pi}{2})\right)} \tag{S.22b}$$

$$I_{rr}(\Theta) = \frac{N}{2\Lambda^2}\pi^2 \tag{S.22c}$$

$$\tag{S.22d}$$

Thus, $\sigma_r^{CRLB}$ becomes :

$$\sigma_r^{CRLB} = \frac{\sigma}{\sqrt{N}\sqrt{\frac{\sigma^2\pi^2}{2\Lambda^2}}} \tag{S.23}$$

We note that in the case of a full modulated/demodulated signal, the signal collected is different and the associated equations must be :

$$N^i(r) = \frac{N}{4}\left(1 + m\cos(\frac{2\pi}{\Lambda}r + i\frac{\pi}{2})\right) \tag{S.24a}$$

$$I_{rr}(\Theta) = \frac{N}{\Lambda^2}4\pi^2 \tag{S.24b}$$

$$\sigma_r^{CRLB} = \frac{\sigma}{\sqrt{N}\sqrt{\frac{4\sigma^2\pi^2}{\Lambda^2}}} \tag{S.24c}$$

The theoretical localization precision of the Gaussian fitting is extracted from [3] :

$$\sigma_{x_c}^{CRLB} = \sqrt{\left(\frac{\sigma^2 + \frac{a^2}{12}}{N}\left(1 + 4\tau + \sqrt{\frac{2\tau}{1+4\tau}}\right)\right)} \tag{S.25}$$

Neglecting the sampling effect of the PSF and the background noise, this parameter can be rewrite simply :

$$\sigma_{x_c}^{CRLB} = \frac{\sigma}{\sqrt{N}} \tag{S.26}$$

The final analytical approximated z Cramér-Rao lower bounds for our experimantal conditions takes the simple form :

$$\sigma_{z}^{CRLB} = \frac{\sigma}{\sqrt{N}}\sqrt{\frac{1}{\sin^2(\theta)}\left(\frac{1}{\frac{\sigma^2\pi^2}{2\Lambda^2}} + \cos^2(\theta)\right)} \tag{S.27}$$

### Supplementary Note 4 : Excitation pattern calibration

Precise knowledge of the different parameters of the excitation pattern is a necessary condition to obtain satisfactory results. More precisely, there are four parameters that need to be characterized exactly to avoid biases for the single fluorophores localization : the orientation angle, the spatial frequency, the tilt angle and the pattern origin. The tilt angle was called $\frac{\theta}{2}$ in the main text in order to simplify calculation and understanding. It is valid if one of the excitation beam is normal to the lateral plane and the other is propagating along the last one. A more general case is developed in this consideration and the tilt angle $\theta$ will now represents the angle of the fringe propagation from the x axis (see **Supplementary Figure 5**)

#### 1 Calibration of the orientation angle

Because the phase information gives access to the position along the fringe pattern direction, it is essential to know the orientation angle in the transverse plane of the camera. The solution is to reconstruct the fringe pattern by localizing single fluorophores in dSTORM buffer deposited at the coverslip (see **Supplementary Note 5**). Then, we reconstruct the phase image (**Supplementary Figure 5 B**) and the orientation angle is retrieved by cross-correlating an iso-phase line of the pattern with a simulated signal with a variable angle. The angle maximizing the cross-correlation results is considered as the actual orientation angle.

#### 2 Calibration of the spatial frequency

Intensity measurements of the sampled PSF allows one to obtain the position according to the direction of modulation. This position is known modulo the fringe periodicity, which means that it is crucial to know it precisely in order not to introduce any bias. We choose to observe a sparse sample composed of 40-nm diameter fluorescent dark red nanobeads. The signal emitted by a single bead is related to its position in the fringe pattern as shown in **Supplementary Figure 5 B**). We move the sample using a nanometric precise motorized stage (PI-P545) on a distance of about five expected fringe periodicities and an acquisition is performed to obtain at least ten sampling points for a period of the modulated signal. By tracking the intensity variation, we reconstruct the signal for the four fringe positions. A sinusoidal fitting of the modulated signals is performed in order to get the spatial frequency of the excitation pattern in the transverse plane with an accuracy below 1 nm by averaging all results for all the beads in the field and also to check the phase difference between the four fringe positions (**Supplementary Figure 5 A**). The fringe periodicity resulting correspond to the projection of the fringe pattern in the lateral plane. The tilt angle of the fringe pattern is necessary to know the effective spatial frequency.

#### 3 Calibration of the tilt angle

3D imaging relies on the use of a tilted fringe pattern where the z position is extracted from the measured phase and the transverse position. Finally, the z information is given by the following formula (**Supplementary Figure 5 B**) :

$$z = \frac{\Lambda\Phi}{2\pi\sin(\theta)} - \frac{x_c - x_0}{\tan(\theta)} \tag{S.28}$$

where $\theta$ is the tilt angle that needs to be calibrated, $x_c$ is the nanometric position obtained by Gaussian fitting, $\Phi$ the phase measured by z-ModLoc, $\Lambda$ the spatial frequency and $x_0$ the origin of the excitation pattern. The first part of the formula (S.28) represents the projection of the fringe pattern along the z axis (see **Supplementary Figure 5 B**). This projection is chosen to be equal to the depth of field in order to avoid any z wrapping. The $\theta$ tilt angle is simply the sum of the two beams propagation tilt angles. By imposing experimental constraints for beam propagation tilt angles and z fringe pattern projection, the tilt angle can be optimized using the z CRLB information (see **Supplementary Note 3**, about 40.1°for a fringe projection of 1 micron). We prepare labeled microspheres as described in the **Supplementary Note 5** to calibrate $\theta$. The measurement of the sphere radius, as well as the center position, gives the theoretical z value with the sphere equation :

$$z = R - \sqrt{(R^2 - (x - x_s)^2 - (y - y_s)^2)} \tag{S.29}$$

Where R is the sphere radius while $x_s$ and $y_s$ are the position of the sphere center. Lastly, we obtain the tilt angle $\theta$ by fitting z obtained by (S.28) to the expected z value obtained by (S.29) and we obtain the curve described in **Figure 2e**, which evidences the performances in terms of accuracy.

#### 4 Calibration of the pattern origin position

The pattern origin position in the field is essential for z-ModLoc imaging as described in the previous paragraph. Any error on $x_0$ could lead to a shift on the z value measured for a single probe. Even if our proposed method is not absolute but rather relative, a z shift value can introduce a z wrapping. Therefore, the position of the pattern origin has an importance for the 3D reconstruction. The pattern origin is corrected so as to center the z histogram at half of the range detection given by the fringe pattern projection along the z axis.

#### Supplementary Note 5 : Localization precision measurement and calibration samples

To obtain the localization precisions displayed in **Figure 2**, we prepared a sample of 40-nm diameter dark red fluorescent beads (10720, Thermo Fisher) randomly distributed on fixed and non-labeled COS7 cells. This operation allow us to obtain a sparse sample with isolated fluorescent nanobeads at different randomly heights. This sample also allows us to be closer to experimental conditions of 3D single molecule imaging.

We added 25 $\mu l$ of a $10^{-5}$ diluted stock solution in 1 ml of Phosphate Buffer Saline (PBS) and we wait 15 minutes before observation for the beads to deposit. Using a custom-made clustering algorithm, we assessed the mean positions of fluorescent isolated nanobeads over more than 50 frames and we estimated the localization precision as the standard deviation of the spatial distribution for each cluster. The calibration of the tilt angle described in **Supplementary Note** 4 was performed by using coated biotin microspheres (Kisker Biotech, PC-B-3.0) labeled with AF647-steptravidin conjugated fluorophores [4]. We prepared a solution containing 500 $\mu l$ of water, 500 $\mu l$ of PBS, 30 $\mu l$ of stock microspheres solution and 1 $\mu l$ of streptavidin-functionalized AF647 (Life Technologies, S21374). The solution was centrifuged during 30 minutes at 10 krpm and the liquid phase was removed and replaced with 100 $\mu l$ of PBS. Then, we vortexed the solution in order to dissolved the deposit. We added 1 ml of PBS to 50 $\mu l$ of the final solution on the coverslip and we waited 1 hour before starting the acquisitions.

To calibrate the excitation pattern, we image single fluorescent fluorophores at the coverslip **Supplementary Note** 4. The sample is obtained by diluting 1 $\mu l$ of stock AF647 florescent probe solution in 500 $\mu l$ of dSTORM buffer (Abbelight dSTORM buffer).

### Supplementary Note 6 : Biological samples preparation

#### 1 Classical biological samples

COS-7 cells were grown in DMEM with 10% FBS, 1% L-glutamin and 1% peni-cillin/streptomycin (Life Technologies) at 37°C and 5% CO2 in a cell culture incubator. Several days later, they could be plated at low confluency on cleaned round 25 mm diameter high resolution# 1.5 glass coverslips (Marienfield, VWR). After 24 hours, the cells were washed three times with PHEM solution (60 mM PIPES, 25 mM HEPES, 5 mM EGTA and 2 mM Mg acetate adjusted to pH 6.9 with 1M KOH) and fixed for 20 min in 4% PFA, 0.02% glutaraldehyde and 0.5% Triton. They were then washed 3 times in PBS (Invitrogen, 003000). Up to this fixation step, all chemical reagents were pre-warmed at 37°C. The cells were post-fixed 10 min with PBS + 0.1% Triton X-100, reduced 10 min with NaBH4, and washed in PBS three times before being blocked 15 min in PBS + 1% BSA. The cells were incubated 1 hour at 37°C with 1 :300 mouse anti-$\alpha$-tubulin antibody (Sigma Aldrich, T6199) in PBS + 1% BSA. This was followed by three washing steps in PBS +1% BSA, incubation during 45 min at 37° C with 1 :300 goat anti-mouse AF647 antibody (Life Technologies,A21237) diluted in PBS 1% BSA and three more washes in PBS. A post-fixation step was performed using PBS with 3.6% formaldehyde during 15 min. The cells were washed in PBS three times and then reduced during 10 min with 50 mM NH4Cl (Sigma Aldrich, 254134), followed by three additional washes in PBS.

#### 2 COS7 cells in collagen matrix

COS-7 cells were grown in DMEM with 10% FBS, 1% L-glutamin and 1% penicillin/streptomycin (Life Technologies) at 37°C and 5% CO2 in a cell culture incubator. Upon reaching confluency, the cells in DMEM were added to 80% collagen, 10% MEM and 5.83% neutralizing solution (RAFT3D Cell Culture Kit, LONZA). The last step was done on ice at 4°C. Then, the cells were plated at $2, 5.10^{-5}$ cells per well on cleaned round 25 mm diameter high resolution #1.5 glass coverslips (Marienfield, VWR). After 30 min at 37°C in the incubator, the exceeding solution was removed and replaced by DMEM without phenol red with 10% FBS, 1% L-glutamin and 1% penicillin/streptomycin. After 24 hours, we did the same steps as described previously.

### Supplementary Note 7 : Data processing workflow

#### 1 Localization on raw data

All processing scripts are based on public available software. First, we remove the background signal on the 512x512-pixel raw frames by subtracting the temporal median of the 10 previous and 10 next frames, pixel per pixel. Then, we detect the PSFs by using a wavelet filtering [5] associated to a low intensity threshold and large spot width filtering parameters. This is necessary due to the large disparities in fluorescence intensity obtained by modulation in the frame. We measure the nanometric position by Gaussian fitting using the Gpufit maximum likelihood estimate Python toolkit [6]. The intensity is then obtained by a photon counting over a 1 $\mu$m x 1 $\mu$m around the position. The second step recombines the four intensities to obtain the phase information for each molecule. The localization array is split in four parts corresponding to the four detection channels. Lateral positions detected in other channels by Gaussian fitting are corrected by using affine transformations in order to correct magnification, rotation and translation caused by misalignment of the detection paths. The top left channel is used as a reference for this operation, and affine transformation parameters are obtained with a simple registration algorithm. We then merge the four localizations corresponding to each fluorophore and apply transmission correction coefficients on each intensity channels as described in the Methods, Optical setup section. Finally, we calculate the phase from the four intensities for each molecule.

For the axial position obtained by ModLoc, we used the x coordinates obtained from the centroid calculation. The z information is obtained from :

$$z = \frac{\Lambda\phi}{2\pi\sin(\theta)} - \frac{x - x_0}{\tan(\theta)} \tag{S.30}$$

#### 2 Drift correction

Drifts are corrected using an axial and lateral custom-made program based on a Direct Cross Correlation (DCC) algorithm [7]. The localization list is divided into several temporal slices based on the frame numbers, the reference slice being chosen as the first and the lateral drift correction is performed at first. The transverse temporal drift is obtained by a 2D cross-correlation of 2 temporal stack. Thus, it is possible to obtain a temporal drift curve whose sampling depends of the size of temporal Stack. When lateral drift are corrected, the operation is repeated along the axial direction. The localizations are binned in 3D images with a voxel size typically set to 60x60x40 nm in x,y,z, and the slice size is generally 500 frames. This operation can be repeated several times by changing the slice size in order to refine the precision.
